# A bi-functional PARP-HDAC inhibitor with activity in Ewing sarcoma

**DOI:** 10.1101/2022.11.10.515994

**Authors:** Louise Ramos, Sarah Truong, Beibei Zhai, Jay Joshi, Fariba Ghaidi, Michael M. Lizardo, Taras Shyp, Hans Adomat, Stephane Le Bihan, Colin Collins, Jeffrey Bacha, Dennis Brown, John Langlands, Wang Shen, Nada Lallous, Poul H. Sorensen, Mads Daugaard

## Abstract

HDAC inhibition has been shown to induce pharmacological BRCAness in cancer cells with proficient DNA repair activity. This provides a rationale for exploring combination treatments with HDAC and PARP inhibition in cancer types that are insensitive to single-agent PARP inhibitors. Here, we report the concept and characterization of a novel bifunctional PARP inhibitor (kt-3283) with dual activity towards PARP1/2 and HDAC enzymes in Ewing sarcoma cells. Compared to the FDA-approved PARP (olaparib) and HDAC (vorinostat) inhibitors, kt-3283 displayed enhanced cytotoxicity in Ewing sarcoma models. The kt-3283-induced cytotoxicity was associated with a strong S and G2/M cell cycle arrest in the nanomolar concentration range and elevated DNA damage as assessed by γH2AX tracking and comet assays. In three-dimensional spheroid models of Ewing sarcoma, kt-3283 showed efficacy in lower concentrations than olaparib and vorinostat and kt-3283 inhibited colonization of Ewing sarcoma cells in an *ex vivo* lung metastasis model. In summary, our data demonstrates the preclinical justification for studying the benefit of dual PARP and HDAC inhibition in the treatment of Ewing sarcoma in a clinical trial and provides proof-of-concept for a bi-functional single-molecule therapeutic strategy.

## Introduction

Poly-(ADP-ribose)-polymerase (PARP) proteins catalyze PARylation of cellular proteins using an ADP (adenosine diphosphate)-ribose subunit of nicotinamide adenine dinucleotide (NAD+) as the donor (1,2). The human genome encodes 17 PARP enzymes where at least PARP1-3 has critical functions in DNA repair, PARP1 being the most characterized (3). PARP1 is essential for the repair of single-strand DNA breaks (SSBs), which is the most common type of breakpoint lesion in cellular DNA (4). When cells encounter SSBs, PARP1 binds the lesion and initiates a PARylation cascade of itself and histones embedded in the chromatin surrounding the SSB lesion. This PARylation event serves as a signal to recruit the SSB repair machinery to patch the lesions before and during DNA replication in the S-phase of the cell cycle (5). Efficient SSB repair is important to prevent replication stress and the more severe double-strand break (DSB) lesions that occur in S-phase when unrepaired SSB lesions collide with the replication forks (4). DSB lesions in S-phase are mainly repaired by homologous recombination (HR) that relies on proteins such as BRCA1 and BRCA2 (6). Deleterious mutations in BRCA1/2 are found in subsets of breast, ovarian, and prostate tumors, and sporadically in other solid tumor indications (7,8). These HR-deficient tumors are indirectly dependent on proficient PARP enzyme activity to avoid accumulation of catastrophic DSBs in S-phase and initiation of cell death (5). This dependency has paved the way for PARP inhibition as a therapeutic strategy to create synthetic lethality in tumor cells with BRCA1/2-deficiencies (9,10).

Currently, there are four approved PARP inhibitors in the clinic: olaparib (approved in 2014), rucaparib (approved in 2016), niraparib (approved in 2017), and talazoparib (approved in 2018). These PARP inhibitors have been widely deployed in cancers with defects in HR DNA repair activity caused by BRCA1/2 mutations (11). Encouraged by the success of PARP inhibitors in BRCA1/2-mutated cancers, research attention has been expanded towards cancer sub-types where HR repair is compromised due to molecular events other than BRCA1/2 mutations. For example, tumors with mutations in RAD51, an enzyme acting downstream of BRCA1/2 in the HR repair pathway, are also sensitive to PARP inhibition (12). This concept is commonly referred to as ‘BRCAness’ and includes all events that mimic BRCA1/2 loss in the context of HR repair (13).

In HR-proficient cancers, the state of BRCAness can be mimicked pharmacologically by inhibition of proteins that impact BRCA1/2 expression. This potentially invites opportunities to broaden the use of PARP inhibitors beyond current clinical practice. For example, impairing dynamic chromatin events related to DNA replication and repair such as histone acetylation can induce pharmacological BRCAness through indirect regulation of HR components (14,15). Recent studies in leukemia, breast cancer, liver cancer, glioblastoma, prostate cancer, and anaplastic thyroid cancer models demonstrated suppression of HR activity with histone deacetylase (HDAC) inhibition that further supports the synergistic potential of HDAC and PARP inhibition (16–23).

Ewing Sarcoma is a highly metastatic bone and soft tissue tumor affecting mainly children and young adults, with a dismal 5-year survival rate of 15-30% for metastatic disease (24). Ewing sarcoma is defined by the presence of specific gene fusion events involving *EWSR1* and the erythroblast transformation specific (ETS) transcription factor *FLI1* (85%) or other ETS-family transcription factors (15%), most often *ERG* (25). These gene fusions encode chimeric oncoproteins (*e.g*., EWS-FLI1 or EWS-ERG) that drive Ewing sarcoma initiation and progression.

Ewing sarcoma cells are sensitive to PARP inhibitors *in vitro* and this sensitivity depends on EWS-FLI1 (26,27). Ewing sarcoma cell line-derived xenografts in mice display sensitivity to FDA-approved PARP inhibitors similar to the responses seen with the standard-of-care chemotherapy temozolomide (26). These observations prompted a phase II single-agent trial in Ewing sarcoma with olaparib, but despite encouraging pre-clinical data, these patients failed to produce durable responses to single agent PARP inhibition (28). The underwhelming response to PARP inhibitors in Ewing sarcoma patients is most likely due to insufficient synthetic lethality and Ewing sarcoma is therefore a prime candidate for exploring pharmacological BRCAness in the context of PARP inhibitors. Here, HDAC inhibition seems attractive based on previously established sensitivity of Ewing sarcoma cells to HDAC inhibitors (29–31). In this study, we have characterized and evaluated kt-3283, a novel dual-specificity single-molecule inhibitor of PARP1/2 and HDAC enzymes in Ewing sarcoma model systems.

## Materials and Methods

### Cell culture

All human cell lines were confirmed free of mycoplasma and maintained at 37°C with 5% CO_2_ and 95% humidity. CHLA10 cells were maintained in Iscove’s Modified Dulbecco’s Medium (Hyclone cat# SH30228.01) containing 1x Insulin-Transferrin-Selenium (ITS) (Thermo Fisher Scientific cat# 41400045) and 20% fetal bovine serum (FBS) (Gibco cat# A3160401). TC32 cells were maintained in RPMI-1640 (Gibco cat# 11875119) with 10% FBS and 1x GlutaMAX supplement (Thermo Fisher Scientific cat# 35050061).

### HDAC activity assay

*In vitro* HDAC activity was measured using the FLUOR DE LYS^®^ HDAC fluorometric activity assay kit (Enzo Life Sciences cat# BML-AK500-0001) following the manufacturer’s protocol. IC_50_ values were calculated using a four-parameter variable slope non-linear regression in GraphPad Prism 8 (GraphPad Software Inc.).

### PARP1 and PARP2 activity assay

*In vitro* PARP1 activity was measured using the HT Universal Colorimetric PARP assay kit (R&D Systems cat# 4677-096-K) and PARP2 activity was measured using the PARP2 colorimetric assay kit (BPS Bioscience cat# 80581) following the manufacturer’s protocol. IC_50_ values were calculated using a four-parameter variable slope non-linear regression in GraphPad Prism 8 (GraphPad Software Inc.).

### PAR formation assay

Cellular PAR formation assays were used to measure the ability of a test compound to inhibit polymerization of PAR. CHLA10 cells were plated on a black, clear-bottom 96-well plate and allowed to attach overnight. Cells were pre-treated with increasing concentrations of test inhibitors for 30 minutes at 37°C before H_2_O_2_ was added to a final concentration of 25 mM and incubated for 5 minutes at RT. After 2x washes with 0.1% Tween-20 in PBS (PBS-T) and 2x washes with PBS, cells were fixed with pre-chilled 70:30 methanol:acetone for 15 minutes at −20°C. Cells were washed with PBS, 2x with 3% BSA in PBS (BSA-PBS) and again with PBS and then blocked with 3% BSA-PBS for 30 minutes at RT. Following 2x washes with PBS and 1x wash with 3% BSA-PBS, cells were incubated for 1 hour at RT with anti-PAR/pADPr monoclonal antibody (R&D Systems cat# 4335-MC-100) diluted 1:250 in 3% BSA-PBS. Plates were washed 2x with 3% BSA-PBS, 1x with PBS, 2x with PBS-T, 2x with PBS and 1x with 3% BSA-PBS, then incubated with goat anti-mouse IgG-FITC (Thermo Scientific cat# F-2761) diluted 1:1000 in 3% BSA-PBS for 1 hour at RT. After washing 2x with 3% BSA-PBS, 1x with PBS, 2x with PBS-T and 3x with PBS, 100 μL PBS per well was added and plates were imaged on an IncuCyte^®^ S3 system (Sartorius). Fluorescence was quantified using the IncuCyte analysis software. Values were normalized to a no primary antibody control and then % PAR formation was calculated by normalizing to a DMSO control. IC_50_ values were then calculated using a four parameter variable slope non-linear regression in GraphPad Prism 8 (GraphPad Software Inc.).

### Cell viability assay

Cells were plated on a 96-well plate (1000-5000 cells per well) in 100 μL appropriate medium and allowed to attach overnight. 100 μL of medium containing dimethylsulfoxide (DMSO) or increasing concentration of test compound was added to each well. Cells were maintained at 37°C with 5% CO_2_ and 95% humidity for 10 days for CHLA10 and 3 days for TC32 and A673.Cell-Titer-Glo^®^ viability assay was carried out for CHLA10.150 μL of media per well was removed and plates were equilibrated at room temperature for 30 minutes, then CellTiter-Glo^®^ assay reagent was added to the wells. The plates were gently shaken on an orbital shaker for 2 minutes and incubated at room temperature for 10 minutes in the dark. Luminescence was measured using a Tecan Infinite M200Pro microplate reader. All measurements were carried out in triplicate. For TC32 and A673, the plates were imaged on an Incucyte S3 live cell imaging system after the treatment period was complete and % confluency was measured using the Incucyte software. Values were normalized to a media-only control and DMSO control to calculate % cell survival. EC_50_ values were calculated using a four-parameter variable slope non-linear regression in GraphPad Prism 8 (GraphPad Software Inc.).

### Cell cycle analysis

Cell cycle arrest profiles were evaluated via propidium iodide staining and flow cytometry. CHLA10 and TC32 cells were plated in 10 cm plates with a cell density of 1.5×10^6^ cells/plate and 2.0×10^6^ cells/plate, respectively. The media was replaced the following day with serum-free media for 24 hours. Cells were treated with olaparib, vorinostat and kt-3283 in a dose escalating manner for 24 hours for TC32 and 48 hours for CHLA10. Combination treatments of olaparib with vorinostat, olaparib with belinostat and olaparib with panobinostat were evaluated at the highest concentration of kt-3283 for both TC32 and CHLA10. Cells were harvested and 1.0×10^6^ cells for each treatment were fixed in 70% ethanol overnight at - 30°C. The suspension was then washed with cold 1X PBS and stained with a propidium iodide solution in 1X PBS (50 μg/mL PI from 1mg/mL stock solution, 0.1mg/mL RNase A, 0.05% Triton X-100) and incubated at 37°C for 40min. Cells were then washed with 1X PBS, filtered through a 40 μm strainer and resuspended with 500 μL of 1X PBS. Samples were then run on FACS and analyzed in FlowJo v10 relative to the DMSO control.

### Alkaline comet assay

To assess single and double-stranded DNA breaks, cells were plated in a 6-well plate at a density of 1×10^6^ cells/well and allowed to settle overnight. Cell media was replaced with serum-free media for 24 hours, then treated with DMSO or increasing concentration of test compound for 24 hours at 37°C with 5% CO_2_ and 95% humidity. Cells were harvested as instructed in Trevigen’s CometAssay^®^ protocol and combined with molten LMAgarose at a ratio of 1:10 and pipetted onto CometSlides^®^. Cells were placed in the dark for 30 minutes and immersed in lysis solution overnight at 4°C. Slides were immersed in Alkaline Unwinding solution for 1 hour at 4°C in the dark then placed in a gel electrophoresis tray and immersed in Alkaline Electrophoresis solution with an applied voltage of 25V for 30 minutes. Samples were washed with dH_2_O and 70% ethanol before staining with SYBR^®^ Gold. Samples were then viewed using a fluorescence microscope.

Acquired images were analyzed using the OpenComet tool on ImageJ (NIH). Further statistics were performed using GraphPad Prism 8.

### Immunofluorescence

CHLA10 cells were plated in a 24-well plate at a density of 3.5×10^5^ cells/well and allowed to settle overnight. Media was replaced with serum-free media for 24 hours, then cells were treated with DMSO or test compound. After 24 hours, cells were fixed with 4% paraformaldehyde (PFA) and stored at 4°C with 1X PBS. Cells were permeabilized with 0.5% Triton-X in PBS before probing with anti-phospho-histone H2AX (Ser 139) rabbit monoclonal antibody (Cell Signaling Technology cat# 9718) and anti-cyclin A2 (EPR17351) mouse monoclonal antibody (AbCam cat# 181591) overnight at 4°C. Cells were then washed with 1X PBS and probed with goat anti rabbit IgG Alexa Fluor^®^ 488 (Abcam cat# ab150077), and mounted with VECTASHIELD Antifade Mounting medium with DAPI solution onto microscope slides. Cells were then visualized on a confocal microscope (Olympus FV3000).

Acquired images were analyzed by quantifying the foci using ImageJ (NIH). Further statistics were performed using GraphPad Prism 8.

### Western blot analysis

Cells were plated in a 6-well plate to 70-80% confluency. After allowing cells to settle overnight, media was replaced with serum-free media for 24 hours. Cells were treated with DMSO or test compound for 48 hours. Cells were harvested in Radioimmunoprecipitation Assay (RIPA) lysis buffer combined with protease and phosphatase inhibitors. Cells were stored at −80°C before running the western blot analysis. Protein yield was assessed using Pierce™ BCA Protein Assay Kit (Thermofisher cat# 23225) and quantified using a spectrophotometer plate reader (TECAN) at 562 nm. A total of 20 μg of protein extracts were loaded per well in 4-15% Mini-PROTEAN^®^ TGX^TM^ Precast Protein gels (Bio-Rad cat# 4561084). After electrophoresis, proteins were transferred onto a 0.2 μm nitrocellulose membrane. The membrane was blocked with LICOR^®^ Odyssey Blocking Buffer in PBS. After blocking, the membrane was incubated with antiphospho-histone H2AX (Ser 139) rabbit monoclonal antibody (Cell Signaling Technology cat# 2577) and GAPDH (D16H11) XP^®^ rabbit monoclonal antibody (Cell Signaling Technology cat# 5174) at 4°C overnight and then incubated with donkey anti-rabbit IRDye^®^ 800CW secondary antibody (LI-COR cat# 926-32213) for 1 hour at room temperature. The membrane was washed with 1X trisbuffered saline with 1% tween-20 (TBS-T) before being scanned with Odyssey scanner (LI-COR). Band intensities were analyzed using the ImageJ software and plotted using GraphPad Prism 8.

### Spheroid formation assay

CHLA10-tdTomato or TC32-tdTomato cells (2500 per well) were added to a 96-well clear round bottom ultra-low attachment microplate (Corning cat# 7007) and allowed to form spheroids for 24 hours or until they reached 200-300 μm in diameter. Medium containing DMSO or increasing concentration of test compound was added to each well and spheroid growth was monitored for 4 days post-treatment using the IncuCyte^®^ Spheroid Analysis system (Sartorius). Images from 4 days post-treatment and day 0 were analyzed using the IncuCyte^®^ Spheroid Analysis software module. Day 4 values were normalized using a normalization factor from day 0 values and EC_50_ values were calculated using a four-parameter variable slope non-linear regression in GraphPad Prism 8 (GraphPad Software Inc.).

### Pulmonary metastasis assay (PuMA)

Procedures involving mice were approved by local animal care committee, University of British Columbia. EGFP-expressing A673 cells (1 ×10^6^ cells/100 μl saline) were injected tail-vein into 6-8-week-old immune-compromised NSG female mice (Jax Laboratories). After injection, mice were euthanized via isoflurane and CO_2_ asphyxiation according to local animal care standard operating procedures. Lungs were insufflated via gravity perfusion with a pre-warmed (37°C) 1:1 mixture of fully supplemented PneumaCult™-ALI media (STEMCell cat #05001) and 1.2% low-melting point agarose (Lonza) as previously described (32). The pluck (heart & lungs) was carefully removed and placed in ice-cold PBS (supplemented with 1X penicillin/streptomycin) for 20 min to allow for agarose solidification. Small lung slices (~2 mm × 4 mm) were obtained by manual cutting with sterile surgical scissors, and 6 slices per condition were chosen for serial imaging at 0 and 14 days post-injection/treatment. Lung slices were maintained in vitro on gelatine sponges partially submerged in 2 mL of PneumaCult™ media +/- compound in a 6-well plate; media +/- compound was refreshed every 3 days. On the day of imaging, lung slices from each group were transferred to a small 35 mm petri dish with a glass coverslip bottom (IBIDI) to permit aseptic widefield fluorescence imaging. The lung slices were imaged on an inverted Zeiss

Observer.Z1 Colibri microscope using 2.5X objective. Lung tumour burden (percent tumor burden) for a lung slice is calculated as the quotient of the summed area of eGFP lesions and total area of lung slice, multiplied by 100, as previously described (33). ImageJ software was used for image processing. This calculation was performed for all lung slices (n = 5) per experimental group. Average values of percent lung tumour burden per group were compared and analyzed in Graph Prism 8 (GraphPad Software Inc.).

## Results

### A bi-specific small molecule with dual activity against PARP1/2 and HDAC enzymes

Through medicinal chemistry cycles, we developed a small-molecule inhibitor prototype (kt-3283) with dual activity against PARP1/2 and HDACs. *In vitro* activity assay kits were used to determine inhibition of PARP1, PARP2 and HDAC by kt-3283 compared to FDA-approved PARP inhibitor olaparib and HDAC inhibitor vorinostat. A wide concentration range of each compound was used to determine IC_50_ values. kt-3283 had an IC_50_ value of 1.89 μM for inhibition of HDACs while vorinostat was about 40-fold lower at 0.05 μM (**Fig. 1A and Supplementary Fig. S1A**). The PARP1 and PARP2 inhibitory activities of kt-3283 were comparable to olaparib, with IC_50_ values for kt-3283 at 0.338 nM and 2.19 nM, respectively (**Fig. 1B, 1C, Supplementary Fig. S1B and Supplementary Fig. S1C**). To further validate the ability of kt-3283 to inhibit PARP1/2 activity, we used a cellular PAR synthesis assay to determine the level of PAR formation. Comparable to olaparib, we determined an IC_50_ of 1.38 nM for the inhibition of PAR formation in cells treated with kt-3283 (**Fig. 1D and Supplementary Fig. S1D**). These data indicate that kt-3283 is able to inhibit both PARP1/2 and HDAC enzymes.

**Figure 1.**
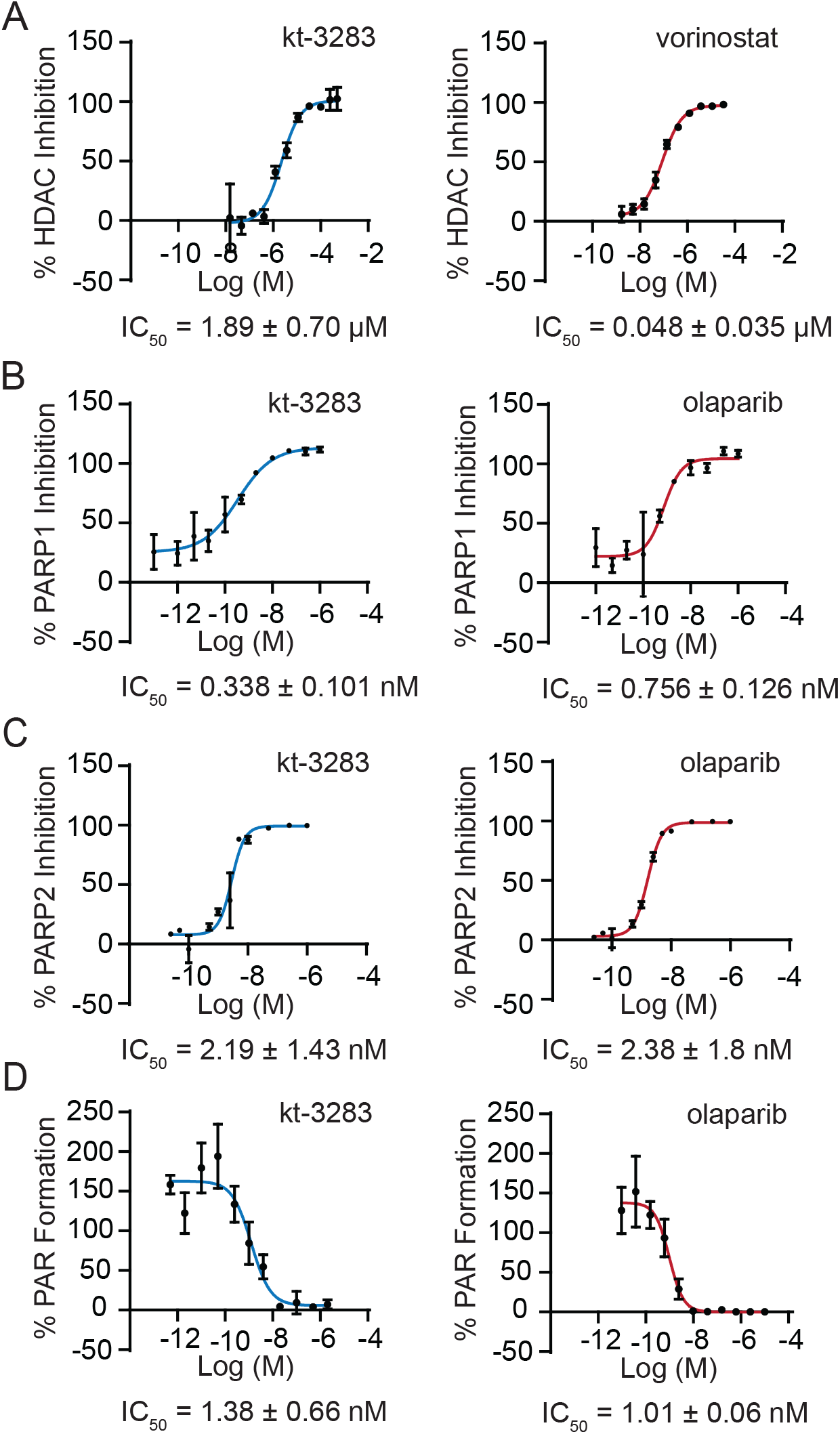
Characterization of a bi-specific small molecule with dual activity against PARP1/2 and HDAC enzymes. **(A)** In vitro HDAC activity in HeLa nuclear extracts treated with kt-3283 or vorinostat. **(B)** PARP1 activity in vitro after treatment with kt-3283 and olaparib. **(C)** PARP2 activity in vitro after treatment with kt-3283 and olaparib. **(D)** PAR formation in CHLA10 Ewing sarcoma cells treated with kt-3283 and olaparib. Values were normalized to control and IC_50_ was calculated as the concentration required to produce 50% inhibition of activity from non-linear regression plots using GraphPad Prism8 software. Data shown are the mean values of n=3 replicates with representative graphs.

### Ewing sarcoma cells are highly sensitive to dual PARP1/2 and HDAC inhibition

In order to investigate the effect of kt-3283 on cell growth, we performed cell viability assays in three Ewing sarcoma cell lines. Using an IncuCyte S3 live cell imaging system to evaluate cell viability, we determined the EC_50_ values after three-day treatment with increasing concentrations of kt-3283, three FDA-approved PARP inhibitors, or three FDA-approved HDAC inhibitors in TC32 and A673 cells. kt-3283 demonstrated higher efficacy in suppression of cell viability than olaparib, niraparib, vorinostat, and belinostat with EC_50_ of 0.0163 μM in TC32 cells (**Fig. 2A and Supplementary Fig. S2A)**. We also detected a similar effect of kt-3283 in A673 cells with a much lower EC_50_ value of 0.0342 μM compared to olaparib, niraparib, vorinostat, or belinostat treatment alone (**Fig. 2B and Supplementary Fig. S2B**). However, treatment with talazoparib or panobinostat showed a more potent inhibitory effect in both cell lines compared with kt-3283 (**Fig. 2A, 2B, Supplementary Fig. S2A and S2B**). To further validate our findings, we also performed CellTiter-Glo^®^ viability assay to determine the EC_50_ values of these tested compounds after 10 days of treatment in CHLA10 cells. Consistent with the IncuCyte assay, the EC_50_ value (0.053 μM) of kt-3283 treatment in CHLA10 cells was also much lower than olaparib, niraparib, vorinostat, and belinostat (**Fig. 2C and Supplementary Fig. S2C**). Taken together, our data demonstrates potent inhibitory effect of kt-3283 in Ewing sarcoma cells compared to most FDA-approved PARP or HDAC inhibitors.

**Figure 2.**
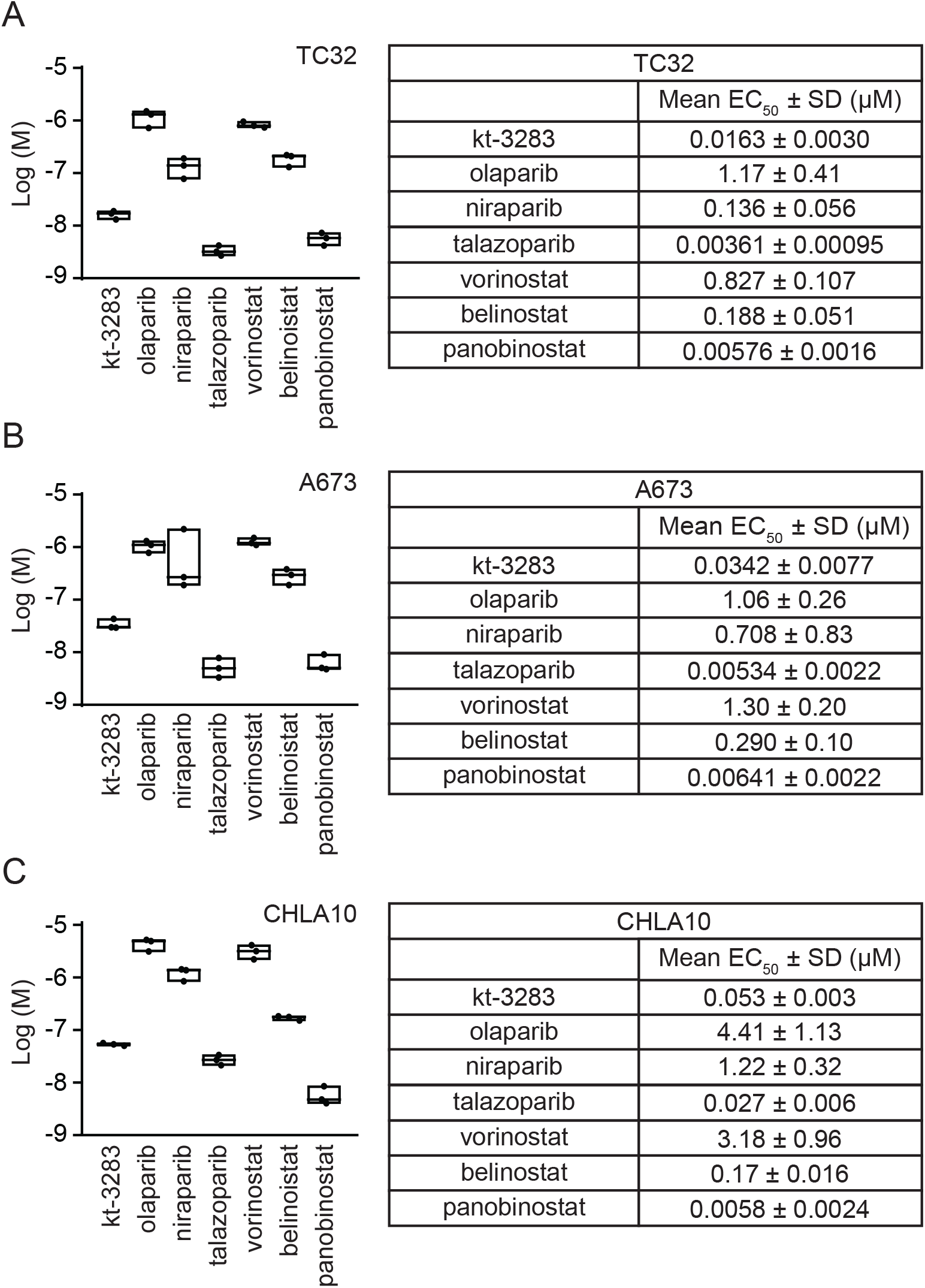
Ewing sarcoma cells are highly sensitive to dual PARP1/2 and HDAC inhibition. **(A)** In TC32 cells, cell viability was examined by IncuCyte S3 live cell imaging system following three-day treatments with increasing concentrations of indicated compounds. EC50 values were calculated as the concentration required for 50% cell viability. n=3 **(B)** EC50 values of tested compounds were also determined in A673 cells using the same experimental condition as in (A). **(C)** EC50 values of indicated inhibitors were determined using CellTiter-Glo^®^ cell viability assays in CHLA10 cells. Cells were exposed to ten-day treatments of increasing concentrations of the inhibitors, and the EC50 values were calculated as the concentration required for 50% cell viability. n=3.

### kt-3283 induces S and G2/M cell cycle arrest in Ewing sarcoma cells

PARP inhibitors and HDAC inhibitors constantly induce S/G2/M and G0/G1 cell cycle arrest, respectively as PARP regulates replication fork progression and HDACs play a major role in regulating the expression of cell cycle checkpoint proteins including cyclin-dependent kinases, cyclin D1 and p21 (34,35). Here we examined the cell cycle profiles of CHLA10 and TC32 cells upon treatment with kt-3283 (3283), olaparib (OLA), and vorinostat (VOR) in both single and combination regimens. Serum-starved cells were treated with increasing concentrations of kt-3283 in complete medium for 24 or 48 h, and displayed strong S and G2/M arrest at and above 0.175 μM in TC32 and 0.25 μM in CHLA10 cells. Similar cell cycle arrest was only observed with olaparib treatment at concentrations higher than 3 μM in TC32 and 14.7 μM in CHLA10, respectively (**Fig. 3A, 3B, Supplementary Fig. S3A and S3B**). Combination treatment with olaparib and vorinostat/belinostat (BEL) at equimolar concentrations of kt-3283 (1 μM and 0.7 μM for CHLA10 and TC32 cells, respectively) had almost no influence on cell cycle phases compared to control (**Fig. 3A, 3B, Supplementary Fig. S3A and S3B**). These data demonstrate kt-3283 has stronger potency to induce S and G2/M cell cycle arrest than olaparib alone or in combination with vorinostat or belinostat in Ewing sarcoma cells.

**Figure 3.**
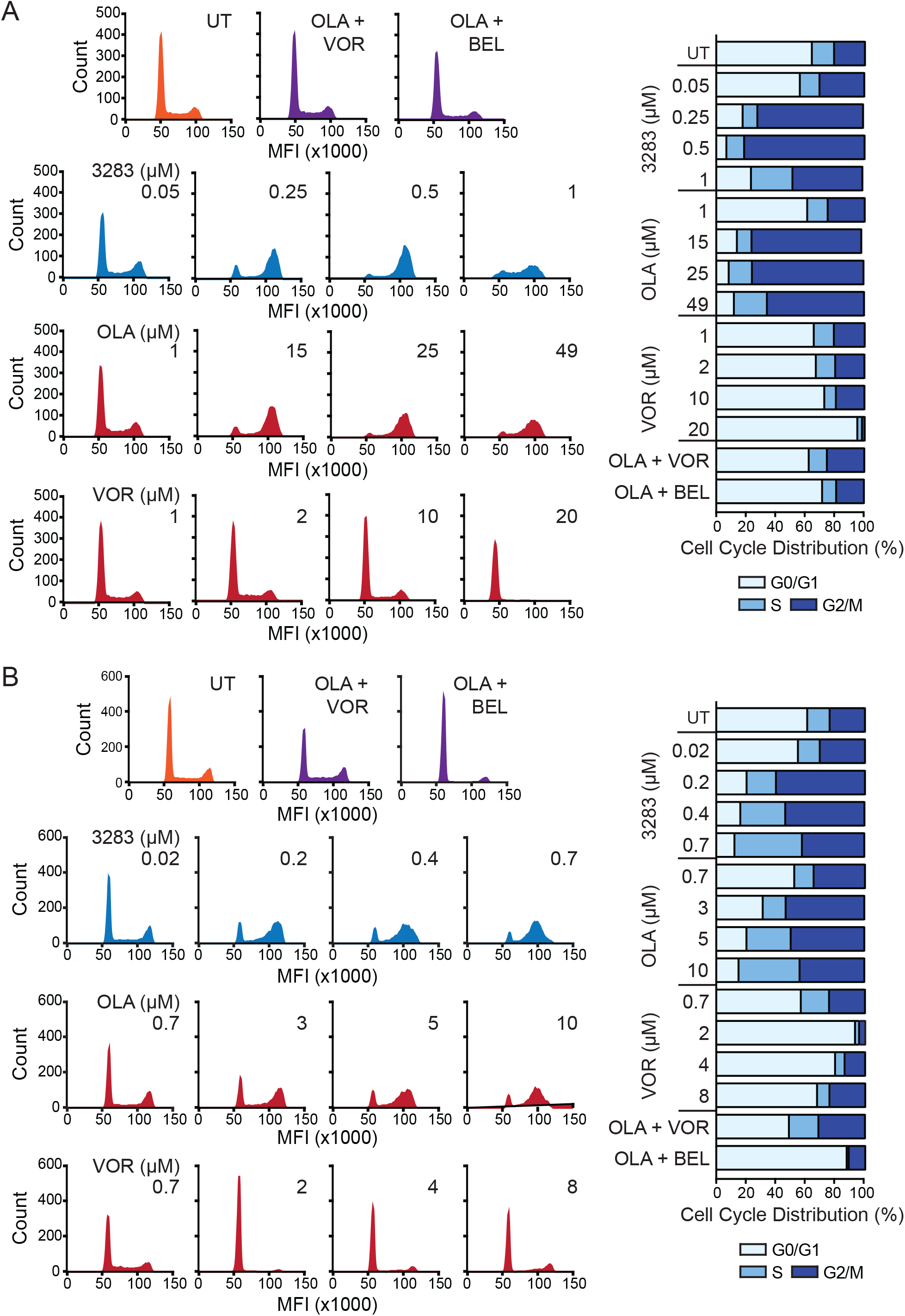
kt-3283 induces S and G2/M cell cycle arrest in Ewing sarcoma cells. **(A)** CHLA10 cells were synchronized at G0/G1 phase by serum starvation for 24 h before treatments with kt-3283, olaparib, or vorinostat as indicated in complete medium for 48 h. Then, cell cycle profiles were examined by propidium iodide (PI) staining followed by flow cytometric analysis as described in “Materials and Methods”. **(B)** Cell cycle analysis was also performed in TC32 cells treated with kt-3283, olaparib, or vorinostat as indicated for 24 h using the same experimental procedures as in (A).

### kt-3283 treatment induces DNA damage in Ewing sarcoma cells

PARP inhibitors and HDAC inhibitors have been reported to induce DNA damage in cells (36). We investigated the effect of kt-3283 on DNA damage compared to olaparib and vorinostat treatment in Ewing sarcoma cells using western blot, immunofluorescence, and comet assay. Phosphorylated histone variant H2AX (γH2AX) is a surrogate marker for DSBs in DNA (37). Dianhydrogalactitol (DAG) was included as a positive control because previous studies in our group showed that DAG induces replication-dependent DNA damage in a variety of cancer cell lines (38,39). Treatment with kt-3283, olaparib or vorinostat induced γH2AX expression in a dose-dependent manner in both CHLA10 and TC32 cells. In comparison to olaparib and vorinostat, kt-3283 was able to induce γH2AX expression at a much lower concentration range (**Fig. 4A, 4B, and Supplementary Fig. S4A and S4B**). Moreover, CHLA10 cells treated with kt-3283 or olaparib also showed dose-dependent γH2AX foci formation in immunofluorescence followed by confocal microscopy imaging with kt-3283 at a much lower concentration range (**Fig. 4C, 4D, Supplementary Fig. S4C and S4D)**. However, vorinostat that induced G0/G1 cell cycle arrest (**Fig. 3A, 3B, Supplementary Fig. S3A, and S3B**) demonstrated milder DNA damage foci formation in CHLA10 cells (**Fig. 4E and Supplementary Fig. S4E**). To further consolidate our data, we also employed alkaline comet assay as it can detect both SSBs and DSBs in cells. There is significant DNA damage in CHLA10 cells treated with 1 μM kt-3283 but not with 1 μM olaparib or vorinostat (**Fig. 4F and Supplementary Fig. S4F).** In summary, our data show kt-3283 is able to induce DNA damage in Ewing sarcoma cells at a much lower concentration range than olaparib or vorinostat.

**Figure 4.**
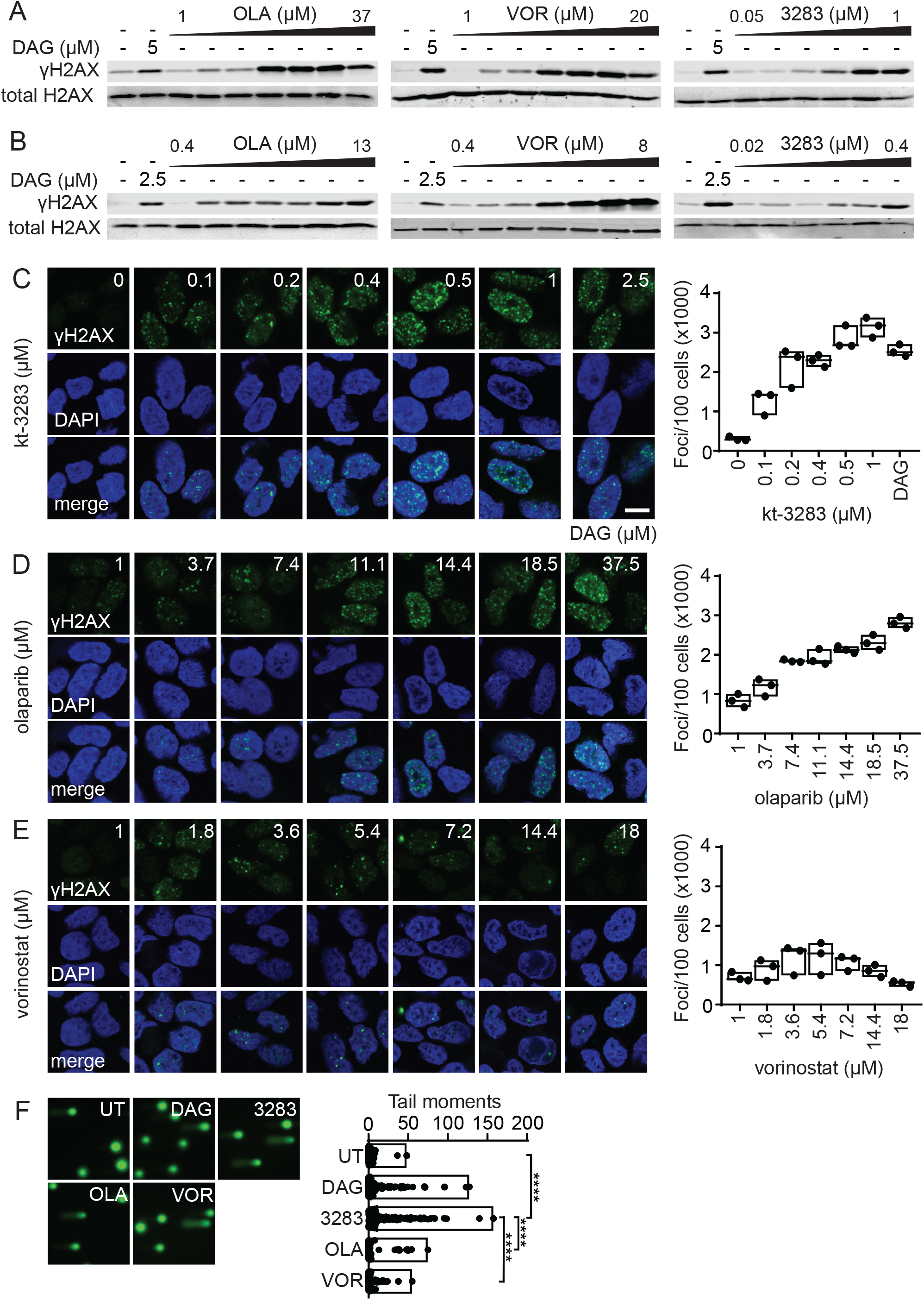
kt-3283 treatment induces DNA damage in Ewing sarcoma cells. **(A)** CHLA10 cells treated with 5 μM dianhydrogalactitol (DAG) or increasing doses of olaparib (1-37 μM), vorinostat (1-20 μM), or kt-3283 (0.05-1 μM) for 48 h and analyzed for γH2AX expression by western blot. **(B)** TC32 cells treated with 2.5 μM DAG or increasing doses of olaparib (0.35-13 μM), vorinostat (0.35-8 μM), or kt-3283 (0.018-0.35 μM) for 48 h and analyzed as in (A). **(C-E)** CHLA10 cells treated with DAG (2.5 μM) or increasing doses of kt-3283 (0-1 μM) **(C)**, olaparib (1-37.5 μM) **(D)**, or vorinostat (1-18 μM) **(E)** for 24 h were analyzed for γH2AX foci by immunofluorescence and confocal microscopy imaging. Scale bar represents 10 μm. **(F)** CHLA10 cells treated with 1 μM kt-3283, olaparib, or vorinostat followed by comet assay as described in “Materials and Methods”. 5 μM DAG was included as positive control. ****p<0.0001

### kt-3283 inhibits 3D spheroid growth and metastasis of Ewing sarcoma cells

Spheroids are three-dimensional (3D) cell aggregates that can mimic tumor behavior more accurately compared to two-dimensional cell cultures (40). To further validate our data, we investigated the effect of kt-3283 on the 3D spheroid model with Ewing sarcoma cells. CHLA10 and TC32 spheroids were established to 200-300 μm followed by treatment with increasing concentrations of kt-3283, olaparib, or vorinostat. The growth of the spheroids was monitored and quantified for four days using the IncuCyte S3 imaging system. The EC_50_ values of kt-3283 in suppression of spheroid growth were much lower than olaparib and vorinostat in both TC32 and CHLA10 cell models (**Fig. 5A and 5B Supplementary Fig. S5A and S5B**). We also examined the effect of kt-3283 on metastatic growth of Ewing sarcoma cells in an *ex vivo* pulmonary metastasis assay (PuMA). Here, colonization of A673 cells in mouse lungs was inhibited by 10 nM or 20 nM of kt-3283 (**Fig. 5C)**. These data suggest that kt-3283 is a potent inhibitor of Ewing sarcoma lung metastasis and that it is more effective than olaparib or vorinostat alone in inhibiting 3D growth in Ewing sarcoma spheroids.

**Figure 5.**
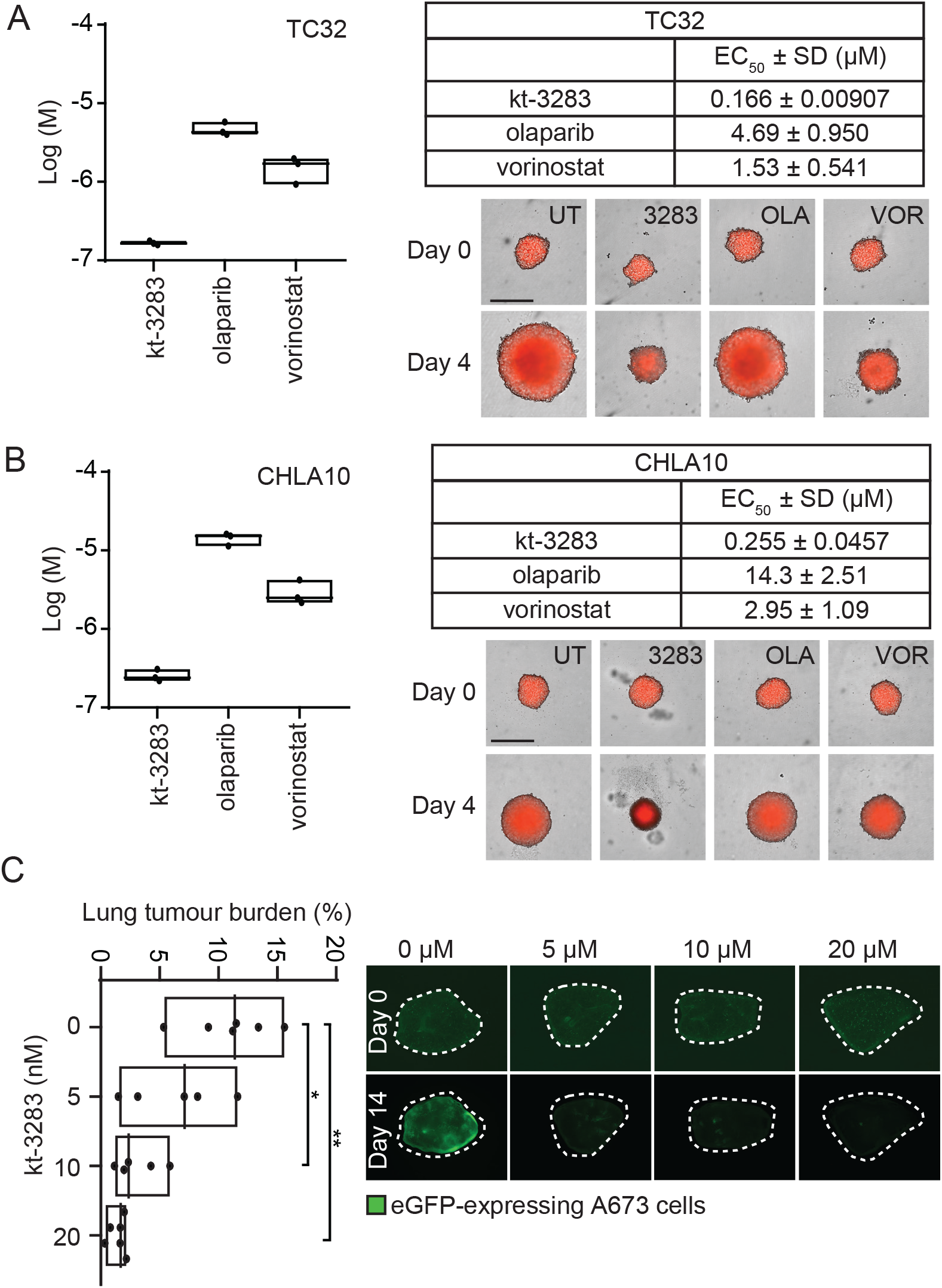
kt-3283 inhibits 3D spheroid growth and metastasis of Ewing sarcoma cells. (A) TC32 spheroid growth following four days of treatment with increasing concentrations of kt-3283, olaparib, or vorinostat was monitored using the IncuCyte^®^ Spheroid Analysis system. The EC50 values were calculated as the concentration required for 50% inhibition of growth from non-linear regression plots using GraphPad Prism8 software. Representative images of TC32 spheroids at day 0 and day 4 with DMSO,I μM kt-3283, 1 μM olaparib, or 1 μM vorinostat are shown with scale bars representing 400 μm. (B) CHLA10 spheroid growth following four-day treatment with increasing concentrations of kt-3283, olaparib, or vorinostat was also examined using the same experimental procedures as mentioned in (A). Representative images of CHLA10 3D spheroids at day 0 and day 4 with DMSO,1 μM kt-3283, 1 μM olaparib, or 1 μM vorinostat are shown with scale bars representing 400 μm. (C) Lung tumour burden following 14 days of treatment with vehicle, 5,10 or 20 nM of kt-3283. n=5. Representative fluorescence images of A673 cells in lung slices following 14 days of treatment with 5 or 10 nM of kt-3283. Scale bar represents 1 mm. *p<0.05 **p<0.

## Discussion

Pharmacological BRCAness can potentially offer a path to take PARP inhibition beyond the BRCA1/2 mutation space and counter potential resistance to PARPi therapy. Epigenetic modifiers such as HDACs, as well as DNA and histone methyltransferases are attractive targets for induced BRCAness in BRCA1/2 proficient cancer scenarios (15,17,19,41–45). At present, four clinical trials using PARPi in combination with the HDACi vorinostat (NCT03259503 and NCT03742245), the DNA methyltransferase inhibitor decitabine (NCT02878785), and the EZH2 histone methyltransferase inhibitor SHR2554 (NCT04355858), are currently ongoing.

FDA has approved three pan-HDACi drugs (vorinostat, belinostat, and Panobinostat) and one HDAC1/2-selective HDACi (romidepsin) for treatment of hematological cancers (46). Histone acetylation attenuates chromatin structure and plays a critical role in recognition and repair of DNA lesions (47). HDACi-induced downregulation of key HR proteins including BRCA1, BRCA2, and RAD51 has been established in a variety of cancers types, and HDACi treatment sensitizes cancer cells to PARPi (15,17–21). This corroborative activity of HDAC and PARP inhibition is particularly interesting in the context of HR-proficient cancer types where PARPi therapy has limited effect on its own, such as Ewing sarcoma. However, dose-limiting toxicity with HDACi therapy is not uncommon in solid tumors (e.g. breast cancer and sarcomas) and has been preventing some therapeutic effects as stand-alone and in treatment combinations (48–50). The HDACi ingredient must be carefully adjusted to prevent over-lapping toxicity events arising from the combination with other therapeutic moieties, which can be challenging when working with different pharmacokinetics profiles.

We have characterized a novel bifunctional PARP-HDAC single-molecule inhibitor, kt-3283, in Ewing sarcoma models to evaluate the potential benefit of combined PARP-HDAC inhibition over stand-alone PARPi or HDACi treatments. kt-3283 has similar PARPi activity as olaparib and slightly lower HDACi activity than vorinostat. However, the dual activity of kt-3283 is 30-to-80 times more cytotoxic to Ewing sarcoma cells than olaparib, and 30-to-60 times more cytotoxic than vorinostat alone. Similarly, kt-3283 induces cell cycle arrest and DNA damage in Ewing sarcoma cells in much lower concentrations than olaparib and vorinostat. When compared in 3D spheroid models, kt-3283 showed efficacy at 30-to-40 times lower concentrations than olaparib and at 5-to-10 times lower concentrations than vorinostat. kt-3283 also hinders metastatic growth of Ewing sarcoma cells in an *ex vivo* pulmonary metastasis model, with a strong inhibitory effect using as little as 10 nM of inhibitor. Combining PARP and HDAC inhibition into one single molecule may also offer a convenient way to prevent resistance to PARPi therapy. For example, Ewing sarcoma and many other solid tumor indications epigenetically suppress expression of the tumor suppressor gene Schlafen 11 (*SLFN11*), which leads to resistance to DNA damage-inducing agents, including PARPi therapy (51–53). Important here, HDACi treatment prompts re-expression of SLFN11 and re-sensitization to PARPi (54–57). In summary, our work provides preclinical justification for studying a novel single-molecule PARP-HDAC inhibitor in Ewing sarcoma with improved cytotoxicity and DNA damage activity as compared to PARPi and HDACi alone. This concept will likely be relevant in cancer indications beyond Ewing sarcoma and potentially offers an opportunity to suppress therapeutic resistance.

## Acknowledgments

This work was supported by a Robert J. Arceci Innovation Award from the St. Baldrick’s Foundation; a St. Baldrick’s Foundation/Martha’s Better Ewing Sarcoma Treatment (BEST) Grant (#663113); an Accelerate grant awarded by the Mathematics of Information Technology and Complex Systems (Mitacs); and Rakovina Therapeutics Inc.

## Data Availability Statement

The data generated in this study are available upon request from the corresponding author.

