## Supplemental Figure S1-S5 for "A bi-functional PARP-HDAC inhibitor with activity in Ewing sarcoma"

Supplementary Figure 1

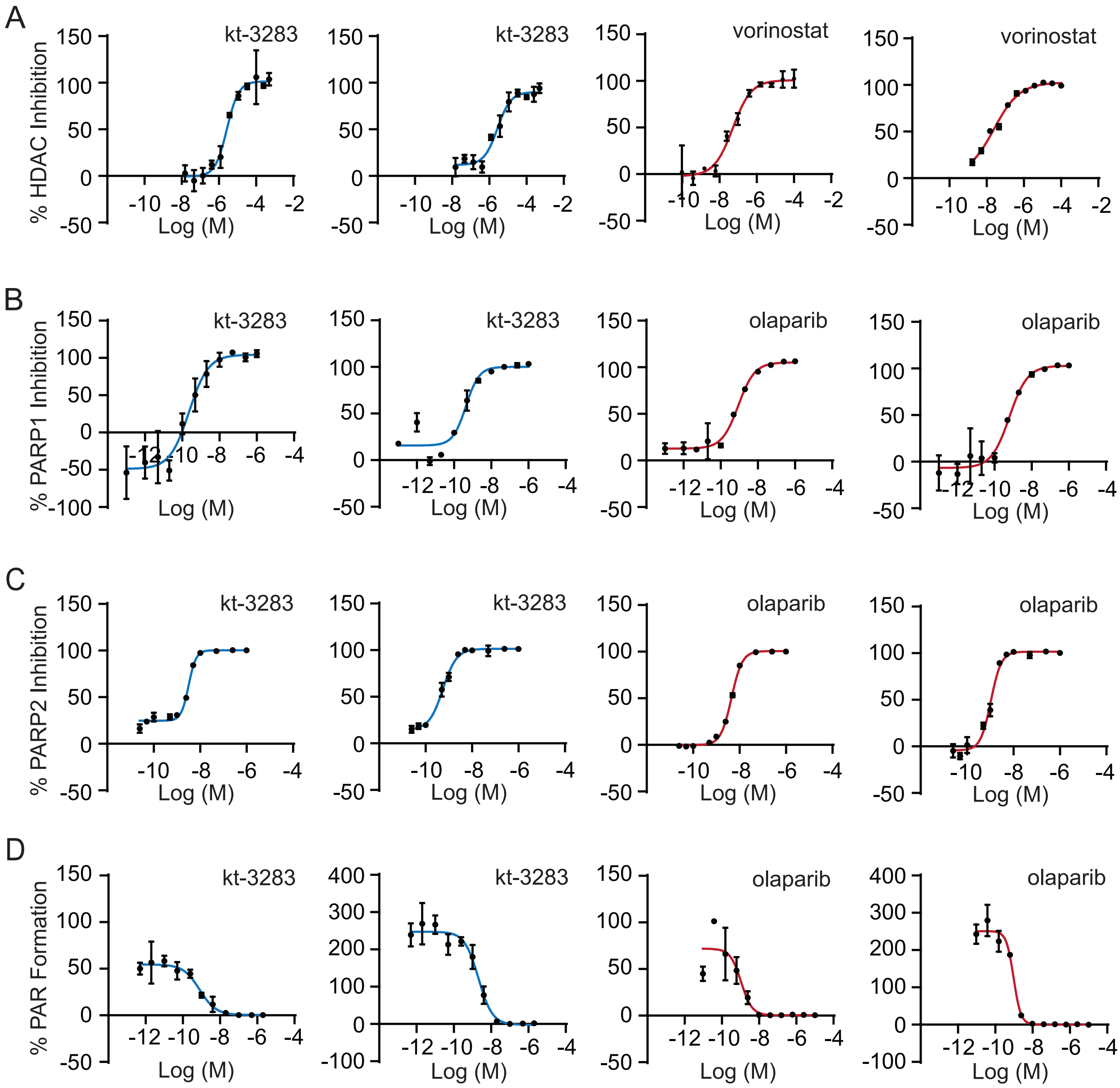


**Supplementary Figure 1**

Replicate graphs of **(A)** In vitro HDAC activity in HeLa nuclear extracts treated with kt-3283 or vorinostat, **(B)** PARP1 activity in vitro after treatment with kt-3283 and olaparib, **(C)** PARP2 activity in vitro after treatment with kt-3283 and olaparib, and **(D)** PAR formationin CHLA10 Ewing sarcoma cells treated with kt-3283 and olaparib. Values were normalized to untreated control.

### Supplementary Figure 2

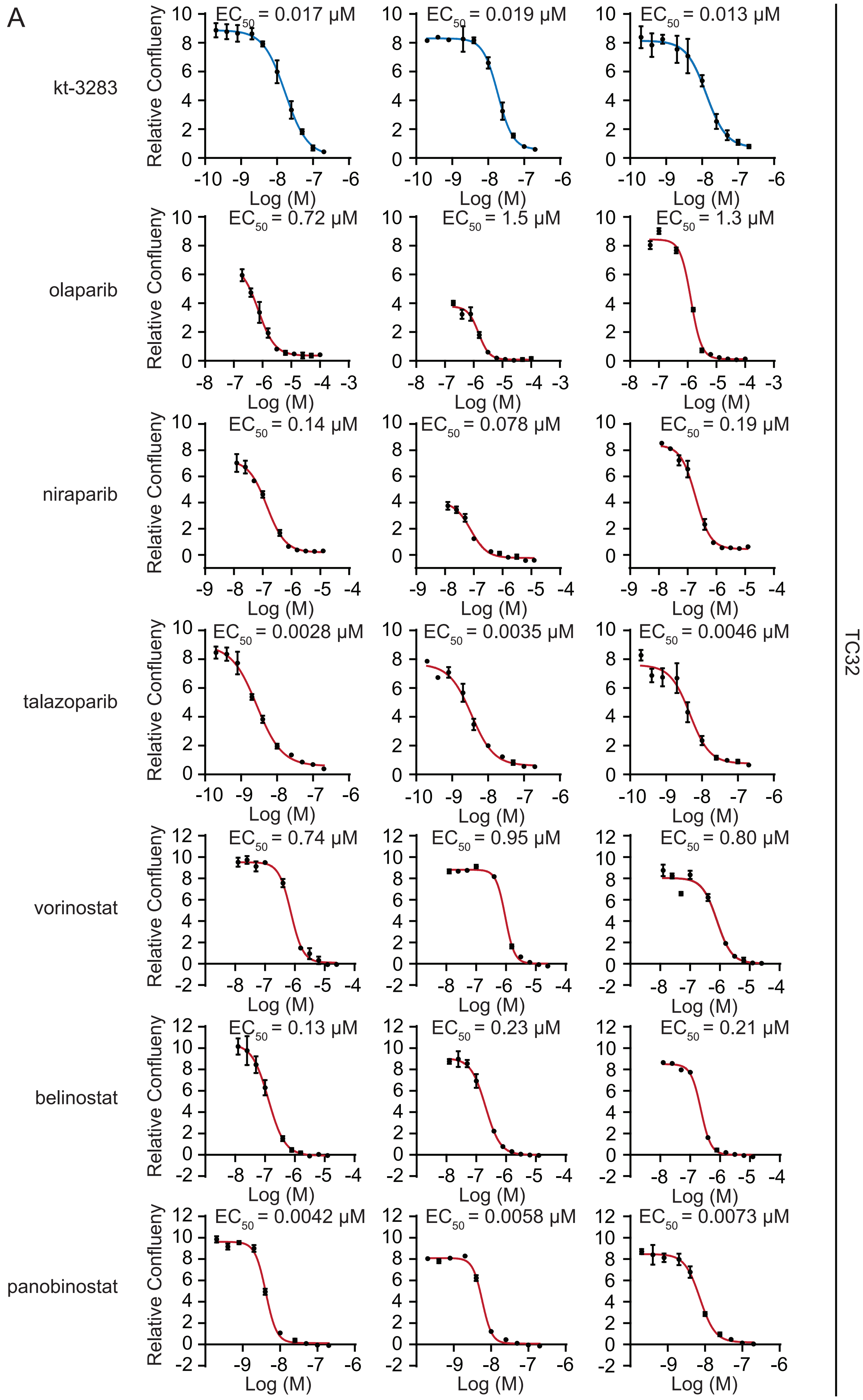

B

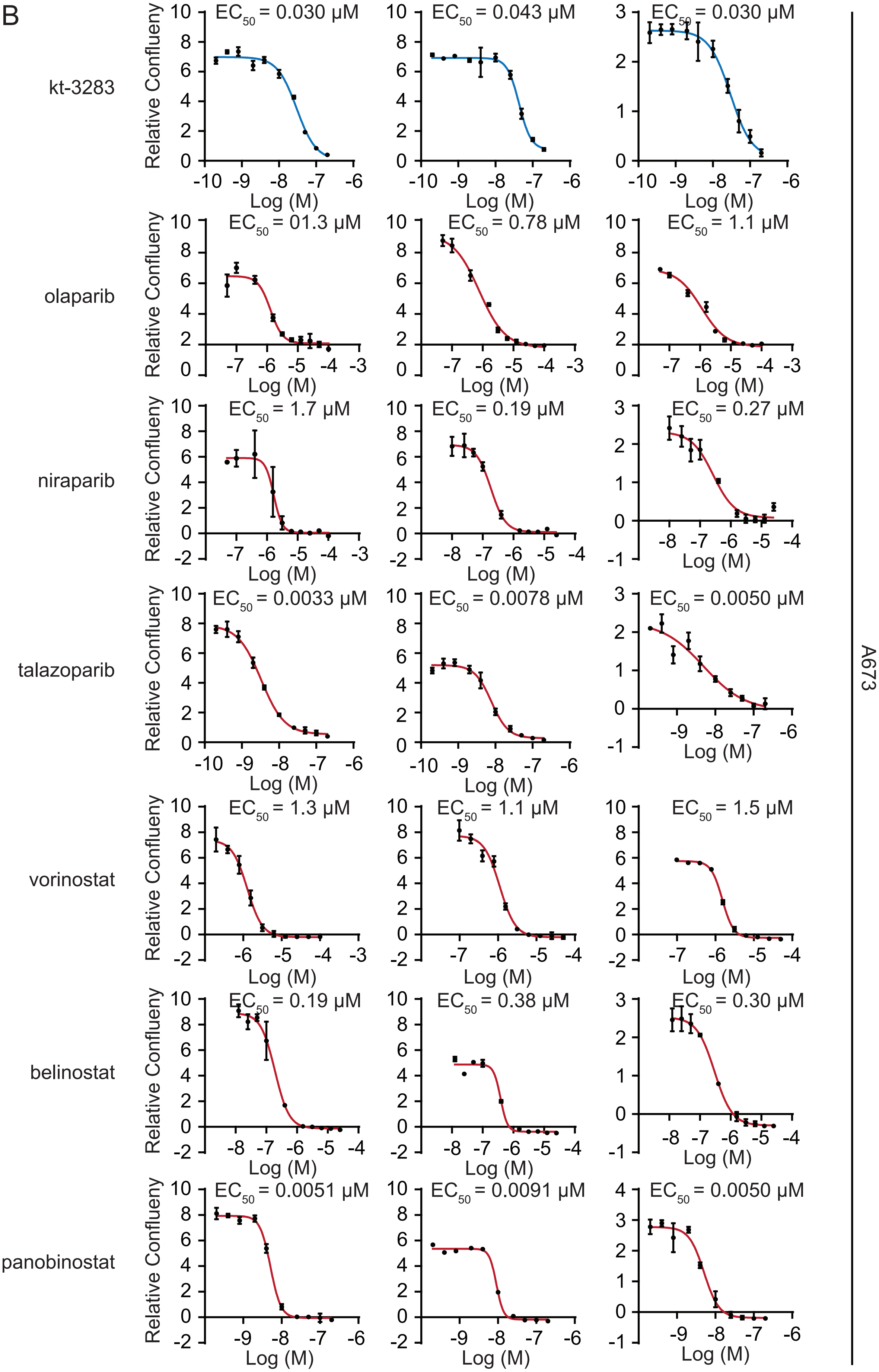
kt-3283
olaparib
niraparib
talazoparib
vorinostat
belinostat
panobinostat
A673
Relative Confflueny
Log (M)
EC50 = 0.030 μM
EC50 = 0.043 μM
EC50 = 0.030 μM
EC50 = 01.3 μM
EC50 = 0.78 μM
EC50 = 1.1 μM
EC50 = 1.7 μM
EC50 = 0.19 μM
EC50 = 0.27 μM
EC50 = 0.0033 μM
EC50 = 0.0078 μM
EC50 = 0.0050 μM
EC50 = 1.3 μM
EC50 = 1.1 μM
EC50 = 1.5 μM
EC50 = 0.19 μM
EC50 = 0.38 μM
EC50 = 0.30 μM
EC50 = 0.0051 μM
EC50 = 0.0091 μM
EC50 = 0.0050 μM

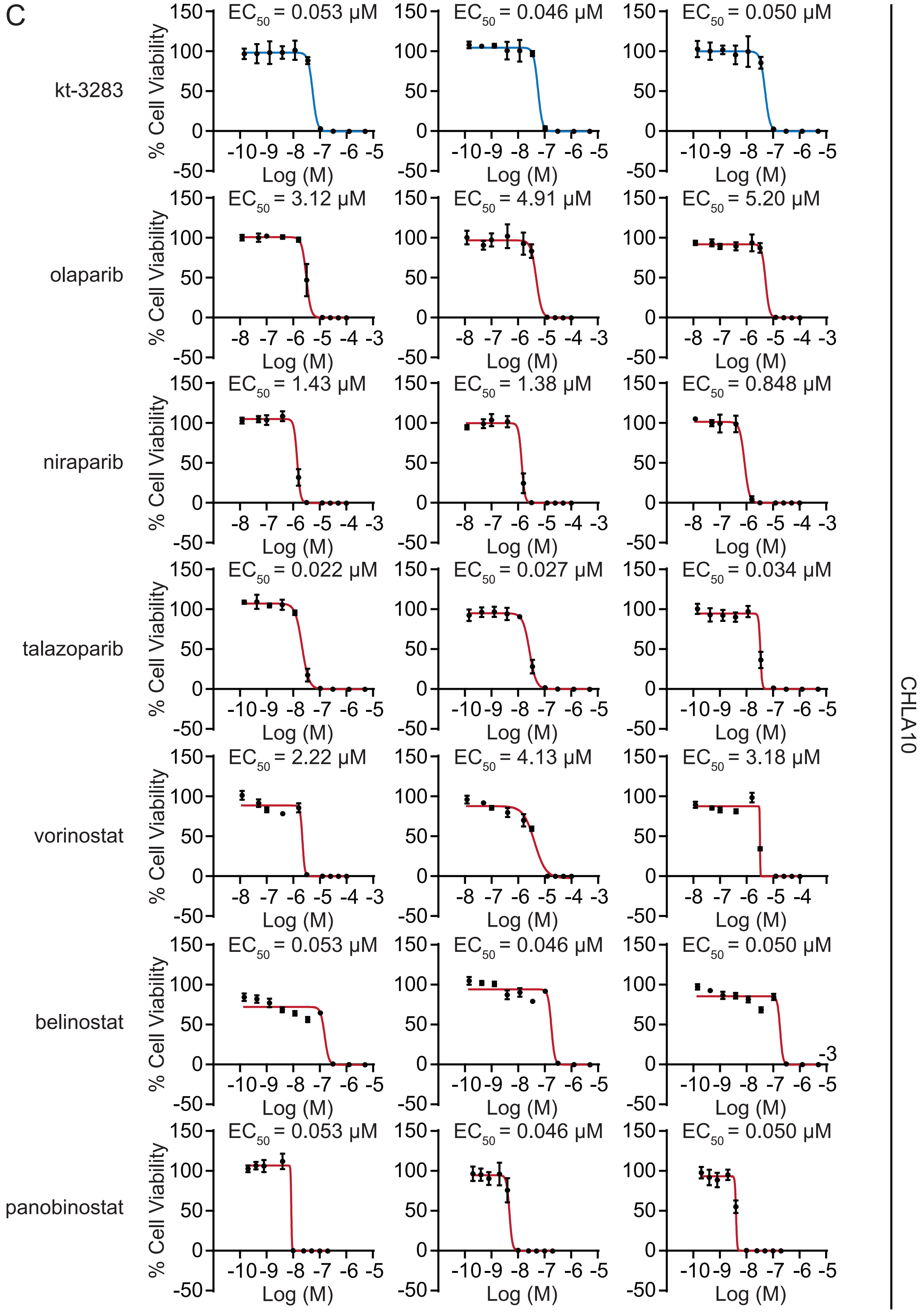


**Supplementary Figure 2**

Replicate graphs of cell confluency following three-day treatments with increasing concentrations of indicated compounds in **(A)** TC32 cells and **(B)** A673 cells. **(C)** Replicate graphs of cell viability following ten-day treatments with increasing concentrations of indicated compounds in CHLA10 cells. Values were normalized to untreated control.

Supplementary Figure 3

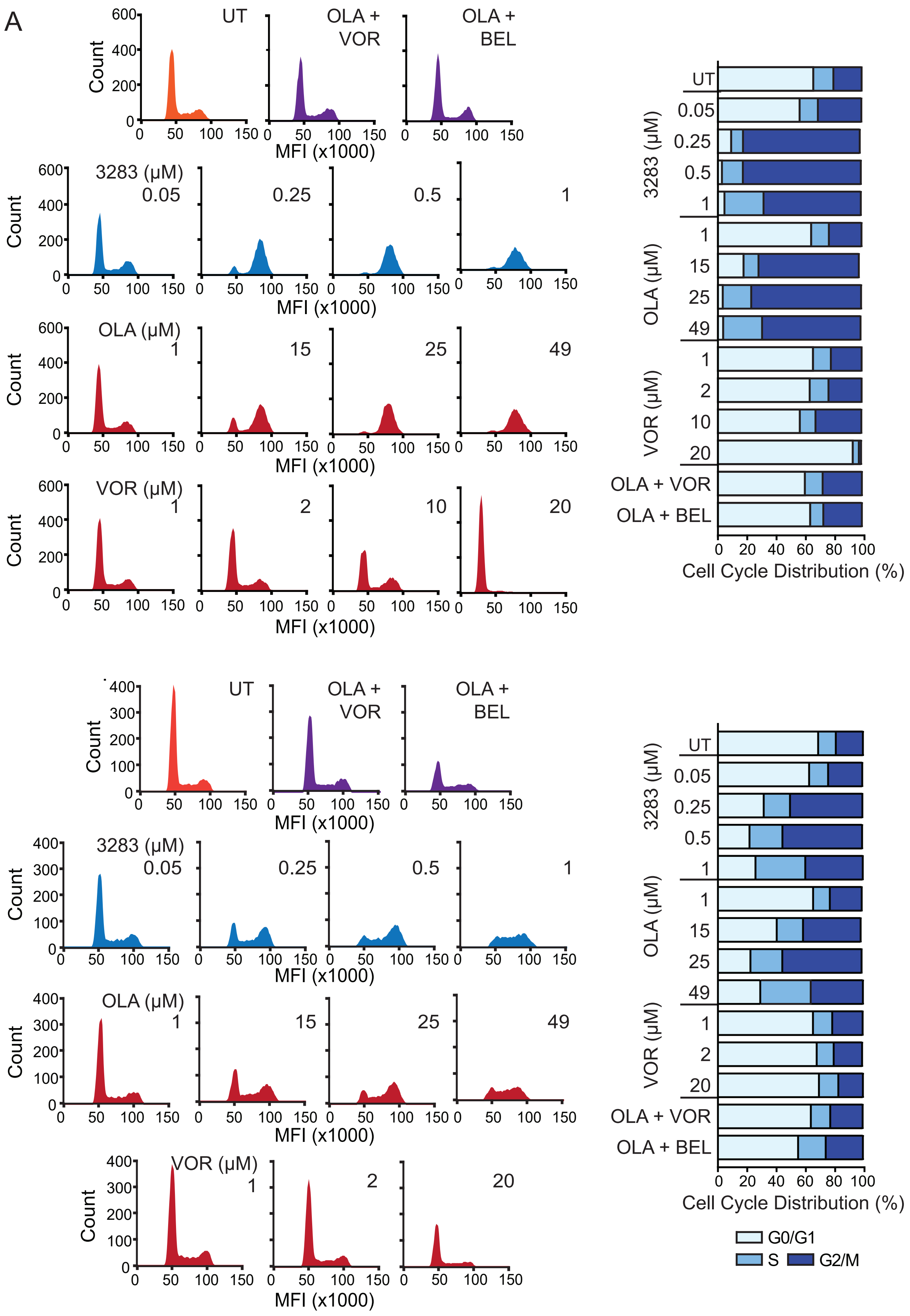

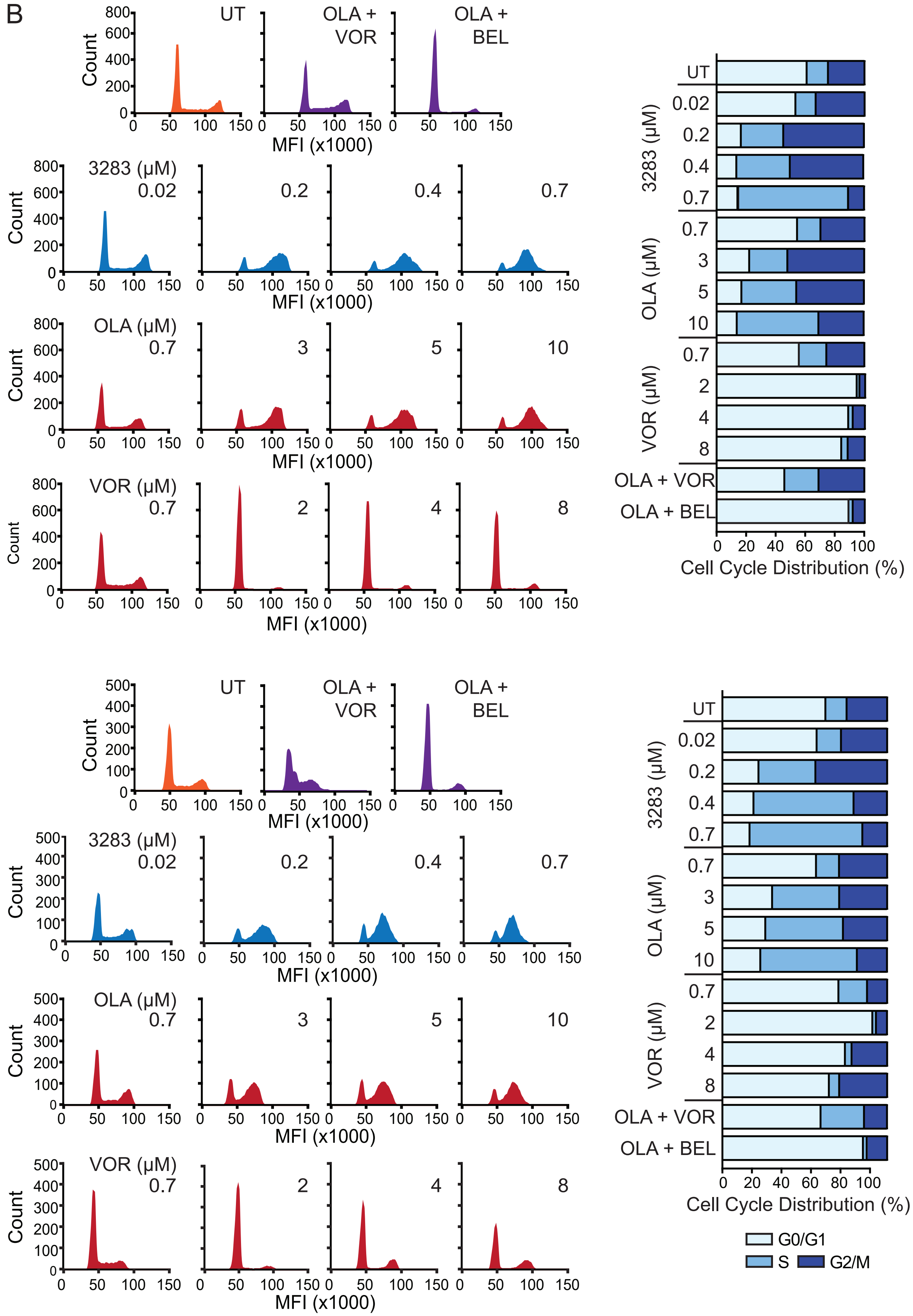


**Supplementary Figure 3**
Replicate cell cycle profiling data for **(A)** CHLA10 cells treated with indicated compounds for 48 h and **(B)** TC32 cells treated with indicated compounds for 24 h.

Supplementary Figure 4

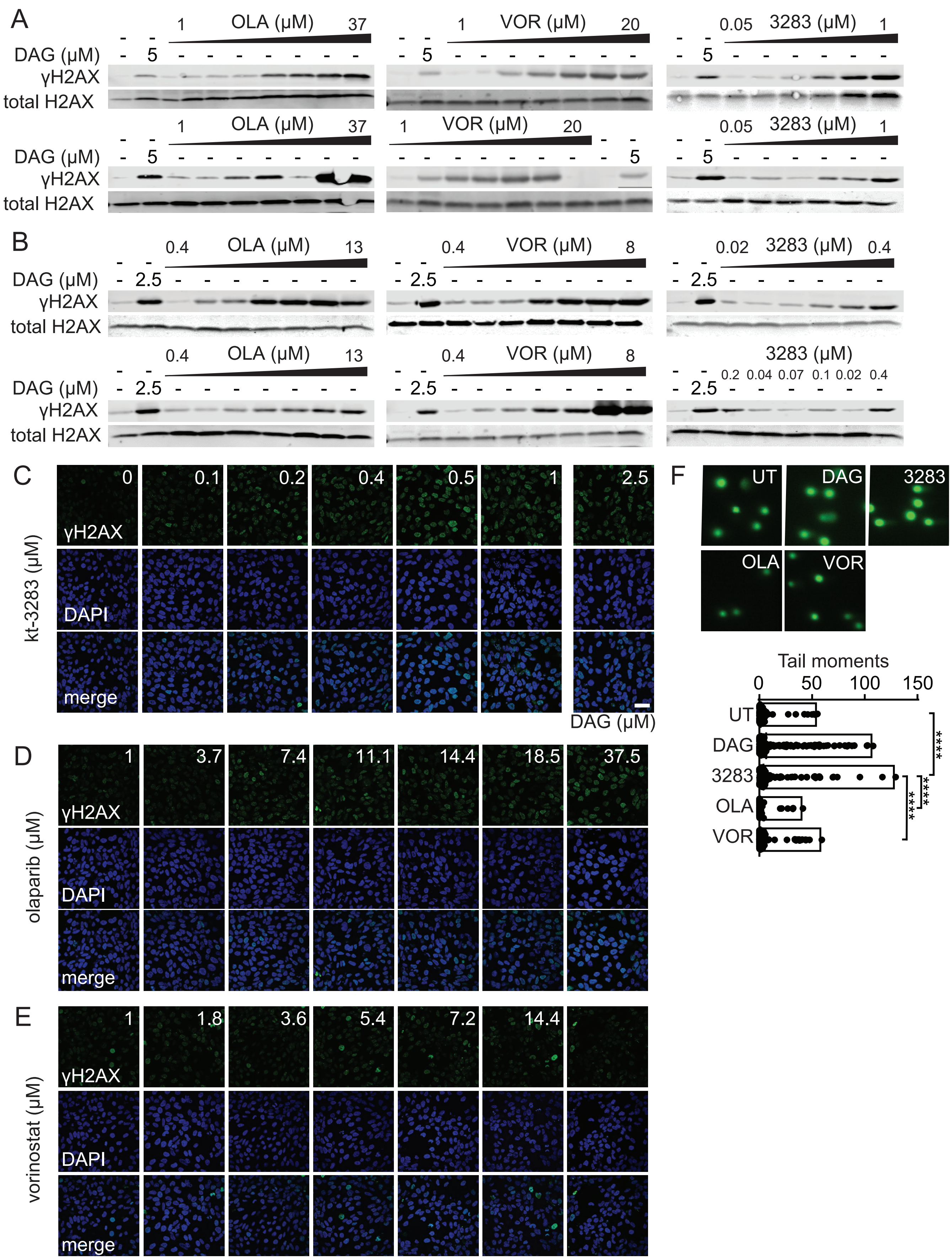

**Supplementary Figure 4**

Replicate western blot images for **(A)** CHLA10 cells and **(B)** TC32 cells treated as indicated for 48 h. **(C-E)** Immunofluorescence images of CHLA10 cells treated as indicated for 24 h. **(F)** TC32 cells treated with 1μM kt-3283, olaparib, or vorinostat followed by comet assay as described in “Materials and Methods”. 2.5 μM DAG was included as positive control. ****$p<0.0001$

### Supplementary Figure 5

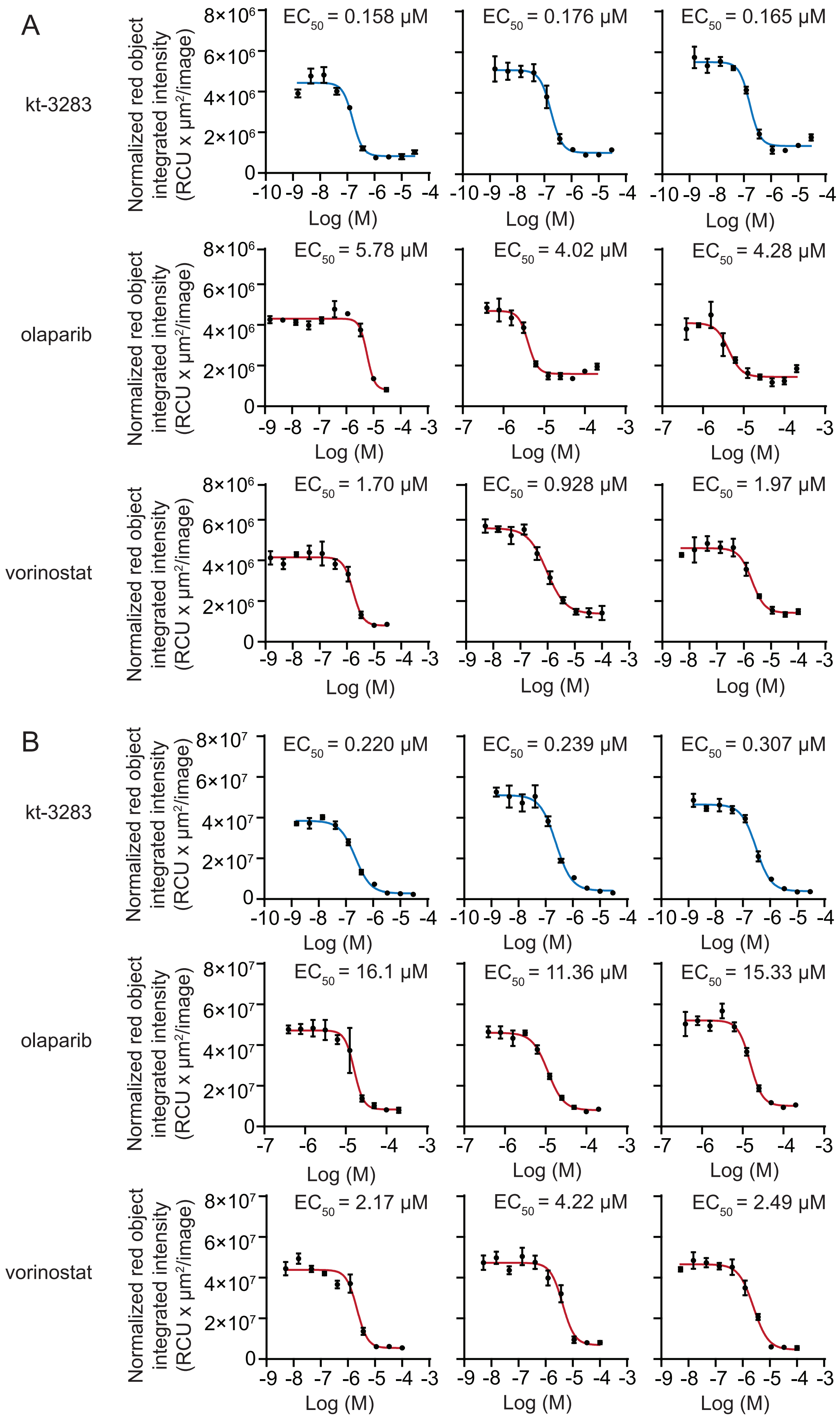

**Supplementary Figure 5**
Replicate graphs of spheroid growth following four days of treatment with increasing concentrations of kt-3283, olaparib, or vorinostat in **(A)** TC32 cells and **(B)** CHLA10 cells. Values were normalized to untreated control.
